# MicNet Toolbox: visualizing and deconstructing a microbial network

**DOI:** 10.1101/2021.11.11.468289

**Authors:** Natalia Favila, David Madrigal-Trejo, Daniel Legorreta, Jazmín Sánchez-Pérez, Laura Espinosa-Asuar, Valeria Souza

## Abstract

Understanding both global and local patterns in the structure and interplay of microbial communities has been a fundamental question in ecological research. In this paper, we present a python toolbox that combines two emerging techniques that have been proposed as useful when analyzing compositional microbial data. On one hand, we introduce a visualization module that incorporates the use of UMAP, a recent dimensionality reduction technique that focuses on local patterns, and HDBSCAN, a clustering technique based on density. On the other hand, we have included a module that runs an enhanced version of the SparCC code, sustaining larger datasets than before, and we couple this with network theory analyses to describe the resulting co-occurrence networks, including several novel analyses, such as structural balance metrics and a proposal to discover the underlying topology of a co-occurrence network. We validated the proposed toolbox on 1) a simple and well described biological network of kombucha, consisting of 48 ASVs, and 2) using simulated community networks with known topologies to show that we are able to discern between network topologies. Finally, we showcase the use of the MicNet toolbox on a large dataset from *Archean Domes*, consisting of more than 2,000 ASVs. Our toolbox is freely available as a github repository (https://github.com/Labevo/MicNetToolbox), and it is accompanied by a web dashboard (http://micnetapplb-1212130533.us-east-1.elb.amazonaws.com) that can be used in a simple and straightforward manner with relative abundance data.

**Author Summary:** Microbial communities are complex systems that cannot be wholly understood when studied by its individual components. Hence, global pattern analyses seem to be a promising complement to highly focused local approaches. Here, we introduce the MicNet toolbox, an open-source collection of several analytical methods for visualizing abundance data and creating co-occurrence networks for further analysis. We include two modules: one for visualization and one for network analysis based on graph theory. Additionally, we introduce an enhanced version of SparCC, a method to estimate correlations for co-occurrence network construction, that is faster and can support larger datasets. We performed method validations using simulated data and a simple biological network. Our toolbox is freely available in a github repository at https://github.com/Labevo/MicNetToolbox, and it is accompanied by a web dashboard that could be easily accessed and manipulated by non-specialist users. With this implementation, we attempt to provide a simple and straightforward way to explore and analyze microbial relative abundance data.

## Introduction

Microbiomes are not a mere collection of independent individuals, but rather, ensembles of intricate constituents, biotic and abiotic, that create highly complex systems where emergent interactions, structures, and functions are crucial for the survival and performance of the whole. The inference of microbial co-occurrence networks may help in understanding emergent properties of these systems [1]: unravelling microbial interactomes [2], evaluating the effects of stress and perturbations in community stability [3], and providing a wide array of novel applications including diagnostics of environmental quality [4] and pathogen identification in disease management [5].

One promising contribution to unravel microbial ecological associations has been the application of network science as an increasingly used alternative to study complex systems [6], whose methods can handle the scale and diversity of high-throughput biological data [1]. Several approaches have been developed to infer microbial ecological associations, such that patterns can be visualized and analyzed within the schematic of a network. One of the most used family of methods is the inference by co-occurrence and correlations, such as: Pearson [7] or Spearman [8] correlation coefficient, Jaccard distance [9] or Bray-Curtis dissimilarity [10], Local Similarity Analysis [11,12], Maximal Information Coefficient [13], MENA that adapts Random Matrix Theory [14,15], SparCC based on Aitchison’s log-ratio analysis [16,17], and CoNet which combines information of several metrics [18,19]. Other types of techniques, such as ordinary differential equations (ODE) models [20] have also been used as an alternative to capture microbial interactions, amongst which the generalized Lotka Volterra equations (gLT) are one of the most used [21] for two-species systems, and potentially useful for three-species systems or larger [22]. Finally, MetaMIS [23], LIMITS [24], and some variations which integrate forward stepwise regressions and bootstrap aggregation [2], offer a different implementation of the gLV equations.

Although the potential value of microbial co-occurrence networks is known, there are several caveats and limitations. To start with, high-throughput genomic data is often associated with low annotation resolution at the species level, which makes it difficult to differentiate between strains and species. This has led to the usage of ASVs (Amplicon Sequence Variants) or OTUs (Operational Taxonomic Units) to obtain a more reliable account of discrete ecological players, leading to hundreds and sometimes thousands of potential organisms [5,25]. However, there are just a few techniques which enable the use of thousands of OTUs/ASVs in the construction of networks for the most diverse ecosystems, such as soil [19]. Furthermore, microbial abundances are normally presented as relative abundance matrices, which creates compositional data sets that are often sparse [1]. Some existing methods are commonly known to provide an efficient approach for compositional effects, spurious correlations, and sparse data handling; nonetheless, biological interaction inferences from compositional data alone should be taken with caution (see **Weiss et al. (2016)** [22], **Dohlman & Shen (2019)** [2], **Hirano & Takemoto (2019)** [26] for a review on performance). Finally, noise or contamination should be expected not be part of the real system captured in the network and the identification of taxa that are not an integral part of a community could be fundamental for more accurate biological interpretations [22].

These issues have led to a high divergence of results when trying to infer direct correlations between OTUs/ASVs [1], therefore algorithms with a more reliable statistical approach are needed. In addition to the intrinsic limitations of inferring interactions from microbial community data, there is a gap in the analysis of networks to obtain the most biological-relevant information: many of the existing methodologies are not easily reachable to the research community, nor do they implement posterior analysis to retrieve information of the co-occurrence network in a clearly and accessible format. Furthermore, at the interpretation level, biologically-meaningful inferences derived co-occurrence networks is still a challenge to be untangled, as signals from co-occurrence may suggest a wide array of phenomena beyond interspecific ecological interactions [1], and most network metrics have debatable or unknown links to relevant concepts in microbial ecology [1,4,27]. In **Table 1**, we present a summary of several network analysis metrics and their current biological interpretation.

**Table 1.**
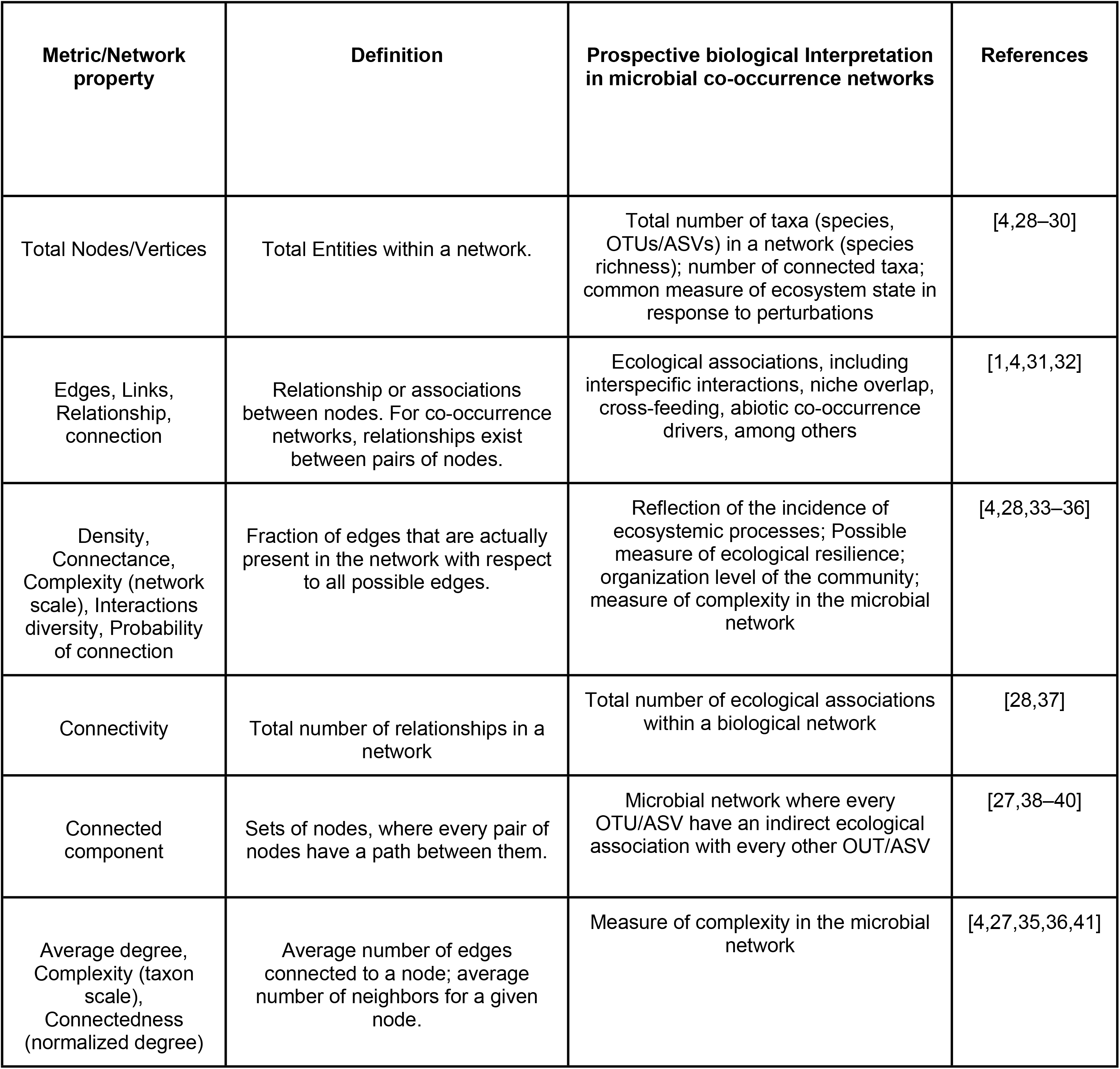
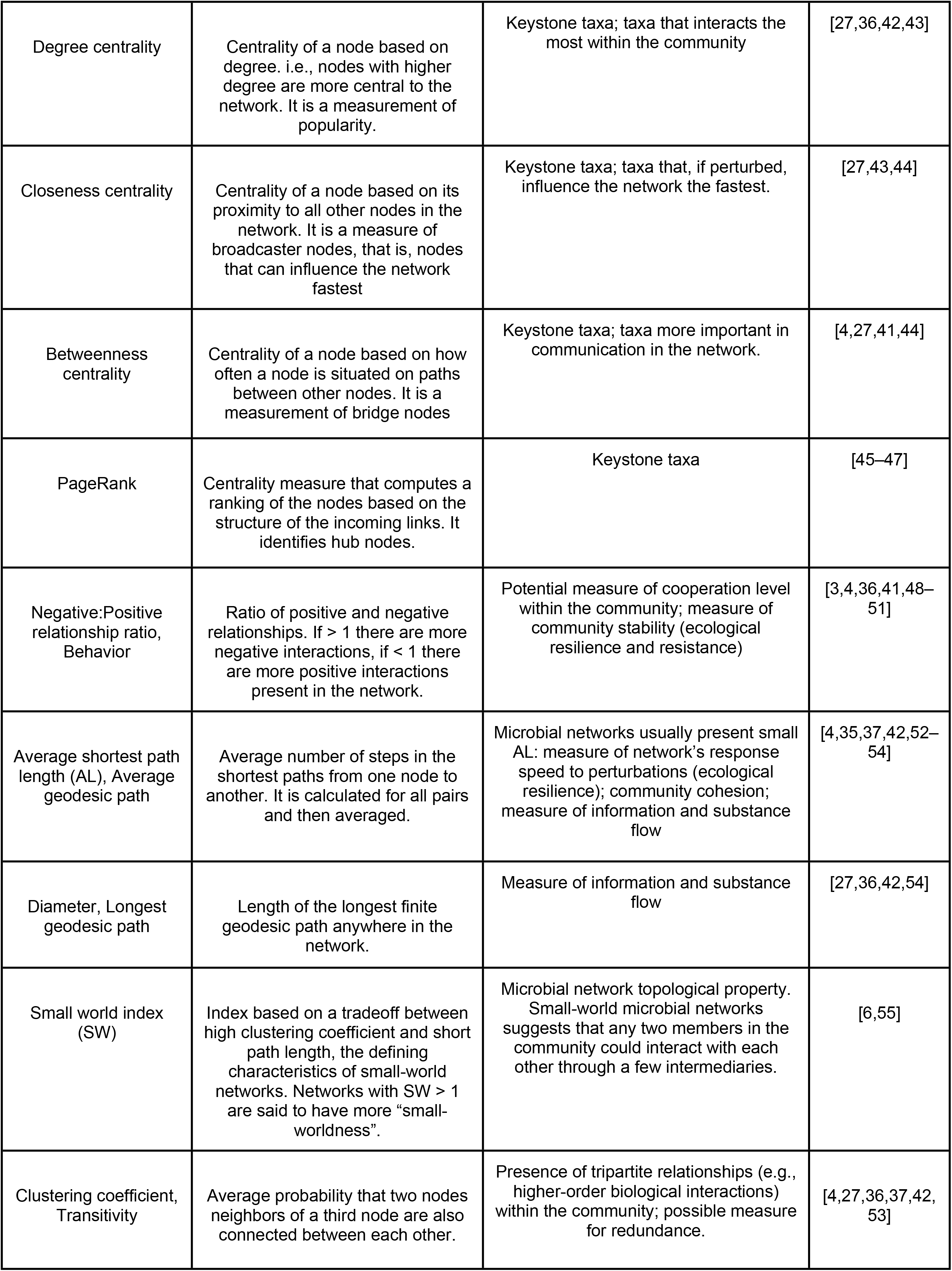
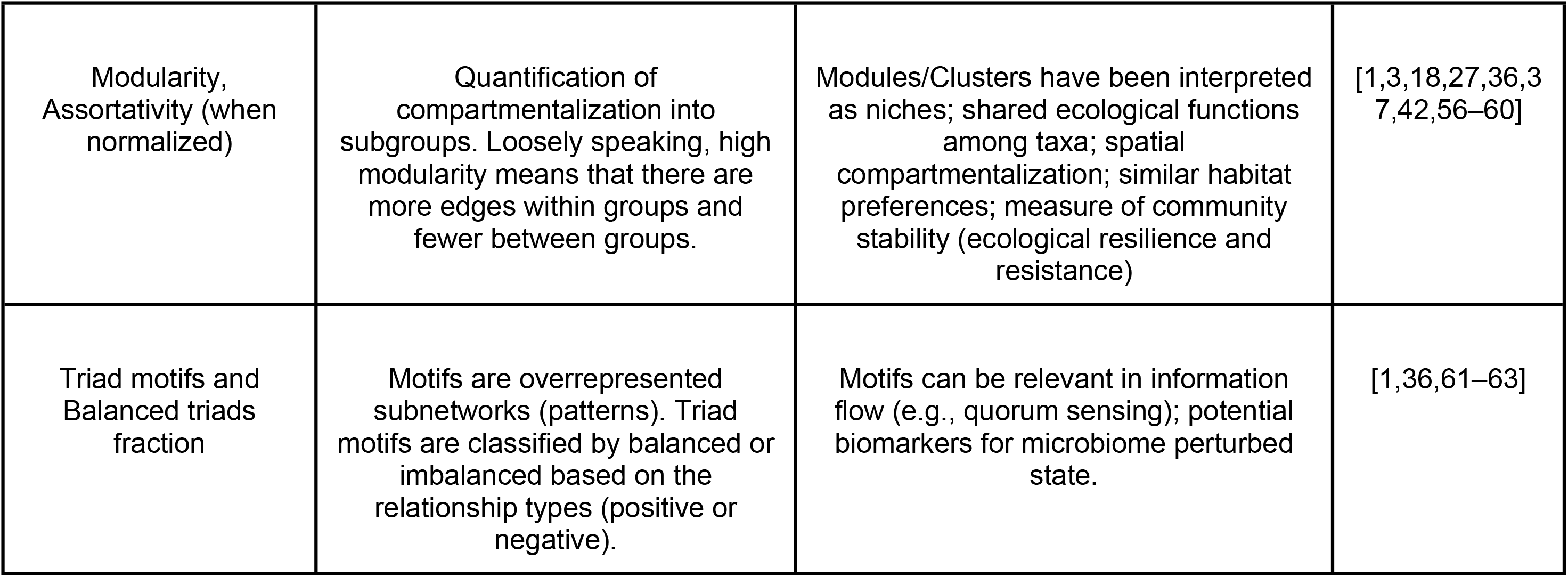
Description of several network metrics and properties currently used in biological networks, including some of their prospective interpretations.

In an attempt to capture the most relevant information from a microbial community in their co-occurrence network and to try to overcome some common issues, we have developed the MicNet toolbox, an open source code to create, analyze and visualize microbial co-occurrence networks. We implemented UMAP [64], a dimension reduction algorithm which has been previously used to identify unique clusters of data in several genomic projects [65,66], given that it is both scalable to massive data and able to cope with high diversity [64]. Moreover, we coupled UMAP with different types of projections and HDBSCAN [67], an unsupervised clustering algorithm, able to identify both local and global relationships, as well as filtering out noise. Finally, we used and enhanced version of SparCC, a compositionally aware algorithm, to infer correlations for network construction [17,22]. Additionally, the toolbox includes several analyses of network theory to inspect the topological properties, robustness, structural balance, communities, and hub nodes that arise in microbial co-occurrence networks. The development of the MicNet toolbox, as an integration of several analyses, attempts to provide an easy-to-use and straightforward implementation towards a comprehensive description of potential local and global patterns for a better understanding of microbial community systems.

## Design and Implementation

### Python Implementation

The code of the MicNet toolbox was built using python 3.9 [68]. The MicNet toolbox uses several standard packages in the Python ecosystem for matrices (pandas v1.3.2 [69,70], numpy v1.20.3 [71], and dask v2021.8.0 [72]), to improve performance (numba v0.53.1 [73]), for temporary storage (h5py v3.2.1 [74]) and to create visualizations (bokeh v2.3.3 [75]). UMAP and HDBSCAN were implemented using packages umap-learn v0.5.1 [64] and hdbscan v0.8.27 [67], whereas network analyses were performed using functions from the networkx v2.6.2 package [76]. In the following sections we explain the different components implemented in the MicNet toolbox which are summarized in **Fig 1**.

**Fig 1.**
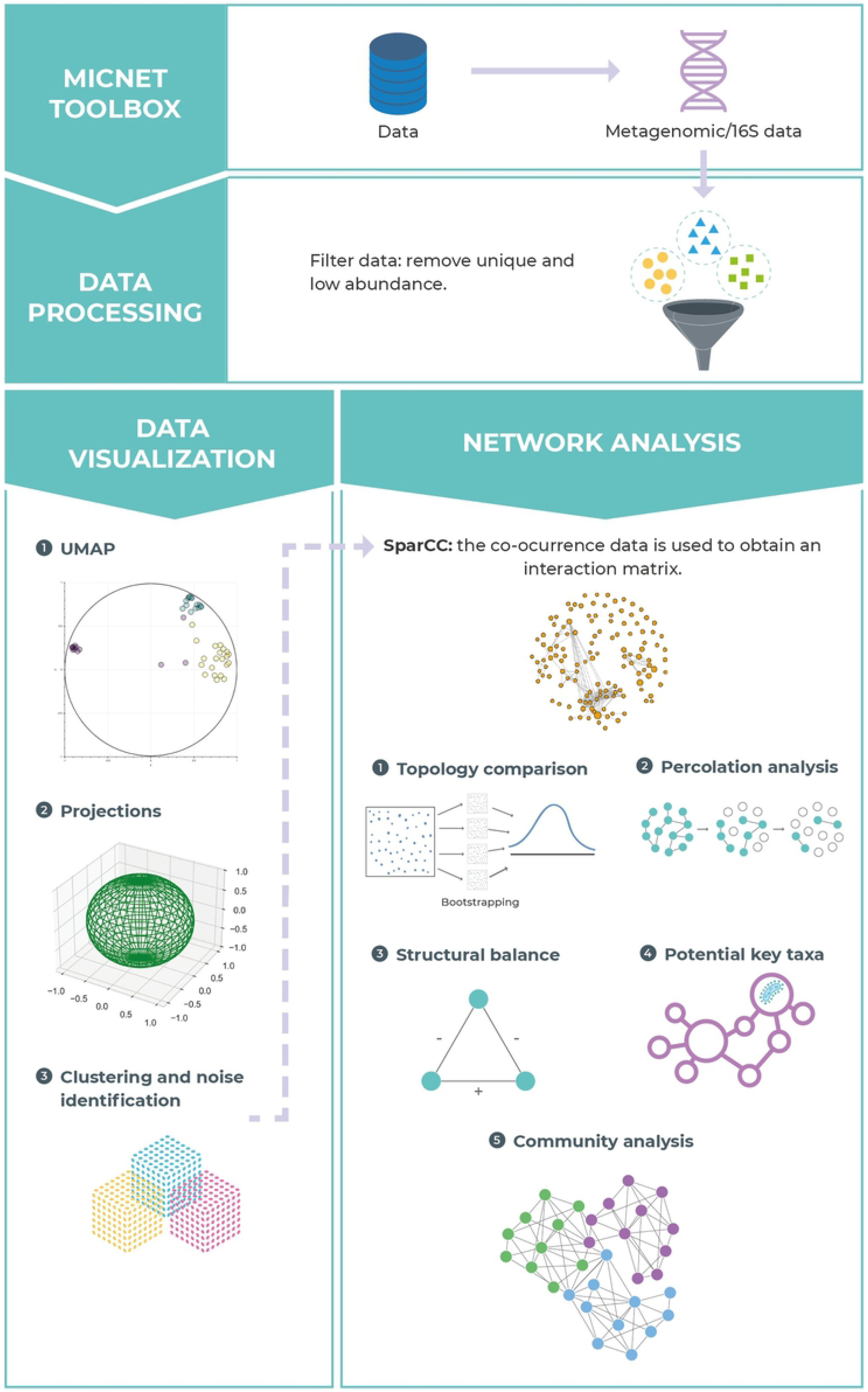
Overview of MicNet toolbox. MicNet Toolbox was designed for visualizing, creating and analyzing microbial networks obtained from compositional data. It includes a data visualization module which uses UMAP and HDBSCAN to find local patterns in the data, and a network analysis module which implements an enhanced version of SparCC and several network analyses such as topology comparison and community analysis, amongst others.

### Input Data

MicNet toobox input data consists of relative abundance/compositional datasets from high-throughput sequencing methods, such as metagenomics or metabarcoding. Prior to building an abundance data table, raw assembled sequences should be OTU/ASV clustered and taxonomically annotated accordingly. MicNet toolbox currently supports abundance data as .tsv files (separated by tabs) or .csv files (separated by commas). In the web dashboard, input abundance data table is filtered by default, removing singletons (< 5 total counts among all samples) and unique (only appearing in one sample) entries. If the user desires, singleton filtering could be deactivated. For SparCC and UMAP/HDBSCAN the first column of the table should contain the OTU/ASV ID and the following columns the abundance data. Taxonomic annotation for the given OUT/ASV can be included in the second column if the user wishes to include taxonomic information in the resulting output. In the case of network analyses, the correlation matrix output by SparCC should be input alongside the UMAP/HDBSCAN output datafile.

### Data Visualization

#### UMAP and HDBSCAN Implementation

A common first step when visualizing high-dimensional data is applying a dimensionality reduction technique. In this toolbox, we implemented UMAP (Uniform Manifold Approximation and Projection), a non-linear dimension reduction technique that favors local data preservation, rather than global data, allowing a better identification of finer scale patterns [64]. We coupled UMAP with HDBSCAN (Hierarchical Density-Based Spatial Clustering of Applications with Noise), a hierarchical clustering algorithm that partitions the data based on their density [67,77]. This clustering technique has been shown to perform well when performed in combination with UMAP dimension reduction [78] and it has been tested with several dataset types, including genetic data, grouping genes into the correct known classes [79]. Thus, we implemented HDBSCAN on the data obtained from UMAP analysis, which represents the abundance data in a reduced space of two dimensions. HDBSCAN analysis not only classifies OTUs/ASVs as belonging to a cluster but also as noise (defined as any point that was not selected in any of the clusters) and outliers (detected with the GLOSH outlier detection algorithm, which works with local outliers).

When running UMAP we set as default a minimum distance of 0.1, number of components of 2, a Helligner output metric and number of neighbors of 15. In the case of HDBSCAN, the default parameters when running MicNet are Bray-Curtis metric, minimum cluster size of 15, minimum sample size of 5 and quantile limit of 0.9 for outlier detection. For our different datasets, the number of neighbors (UMAP), and minimum cluster/sample size (HDBSCAN) parameters were set depending on the input microbiome dataset. We used Bray-Curtis dissimilarity as the distance metric for UMAP, as it is a standard metric for biological datasets. In the web dashboard, UMAP and HDBSCAN parameters can be modified by the user to visualize the results according to each set of microbiome data.

### New Implementation of SparCC

To obtain the correlation matrix from relative abundance data, we implemented a modified version of the SparCC algorithm, a robust approach to discard spurious correlations when dealing with compositional data [17,22]. Although the original SparCC algorithm was not altered, several modifications were made to improve and scale the SparCC estimation matrix. We made three main changes to the original code; first, the code changed from Python version 2.7 to 3.9. Second, we use Numba and Dask in some parts of the matrix processes, namely functions or operations, with two main improvements: parallelization of operations and scalability in the size of the estimated matrices. Finally, the original SparCC version stores each estimation step in RAM, as arrays in NumPy. Although storing in RAM is efficient for small data sets, with large data the required memory increases rapidly depending on the interaction numbers and the size of the dataset. Thus, to overcome this problem, we store each estimation step on disk as hdf5 binary format. These changes made it possible to calculate the SparCC estimates with good performance in easily accessible computing resources. SparCC p-value test on the inferred correlation was not modified, it was calculated with a Monte-Carlo simulation (with default *n* = 50) as done by **Friedman & Alm (2012)** [17] and the default value is to calculate one-sided p-values, although this can be modified by the user.

To set SparCC parameter values, we perform a parameter search in our more complex study case from the communities reported by **Espinosa-Asuar et al. (2021)** [80]. We performed an independent parameter sweep on each parameter, varying the exclusion threshold from 0 to 1 in steps of 0.1, the number of iterations from 10 to 100 in steps of 10 and the exclusion number from 10 to 100 in steps of 10. For each parameter value, we calculated the number of correlations found and selected the parameter value when this number stabilized (**Fig S1: Supplementary material**). The final parameter values used in the databases presented here were: 50 iterations, an exclusion number of 10, exclusion threshold of 0.10 and 100 simulations for p-value calculation. However, this can be modified by the user both in the dashboard and when running the code from the github repository.

### Network Analyses

Network analyses were performed to characterize both the overall structure and the local interactions of the microbial co-occurrence network, in which each OTU/ASV is represented as a node and the correlations found by SparCC as undirected weighted edges, such that an edge between two nodes implies a relationship between the two corresponding OTUs/ASVs. Given that most network analyses can only handle positive interactions, we normalized the SparCC correlation matrix from −1 to 1 to a range from 0 to 1, except for the structural balance analysis which directly uses the positive and negative correlation values. All available network analyses in the MicNet toolbox are described as follows.

#### Network Topology Comparison

Networks have several large-scale structural measurements to characterize their topology. For this purpose, MicNet calculates the following structural metrics: 1) network density, using networkx function nx.density, 2) average degree, calculated as the mean of all nodes degree using numpy mean function, 3) degree standard deviation, using numpy std function, 4) ratio of positive-negative relationships calculated simply as the number of positively weighted edges divided by the number of negatively weighted edges, 5) average shortest path length using nx.average_shortest_path_length, 6) clustering coefficient nx.average_clustering function, 7) modularity, calculated with function nx.modularity, using as network modules those obtained with the nx.greedy_modularity_comminuties algorithm, and 8) the diameter, which was calculated using nx.diameter function. Finally, we have added a custom function that calculates a small-world (SW) index as suggested by **Humphries & Gurney (2008)** [55]. The SW index is calculated as:

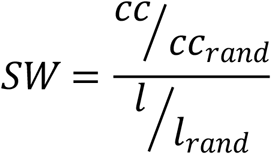

Where *l* and *cc* are the average shortest path length and clustering coefficient of the experimental co-occurrence matrix, respectively. Analogously, *l*_*rand*_ and *cc*_*rand*_ are the average shortest path length and clustering coefficient, accordingly, of a comparable random network with the same number of nodes and density. The random network was built using the function nx.erdos_renyi_graph. This is done several times, with default *n* = 50, and the mean value of SW is returned.

MicNet includes the computation of the distribution of several of this large-scale metrics under the assumption that the underlying topology is: 1) a random Erdos-Renyi network [81] built using function nx.erdos_renyi_graph, 2) a small world Watts-Strogatz network [82] built using nx.watts_strogatz_graph function, or 3) a scale-free Barabási-Albert network [83] built using nx.barabasi_albert_graph function. A short description of these canonical topologies can be found in **Fig 2**. This allows the comparison of the query data against the different topologies. These simulated networks are built with the same number of nodes, density and average degree as the experimental data, and correlations are drawn from a uniform distribution from −1 to 1. Finally, the simulated networks are made symmetrical to be comparable with the SparCC output correlation matrix.

**Fig 2.**
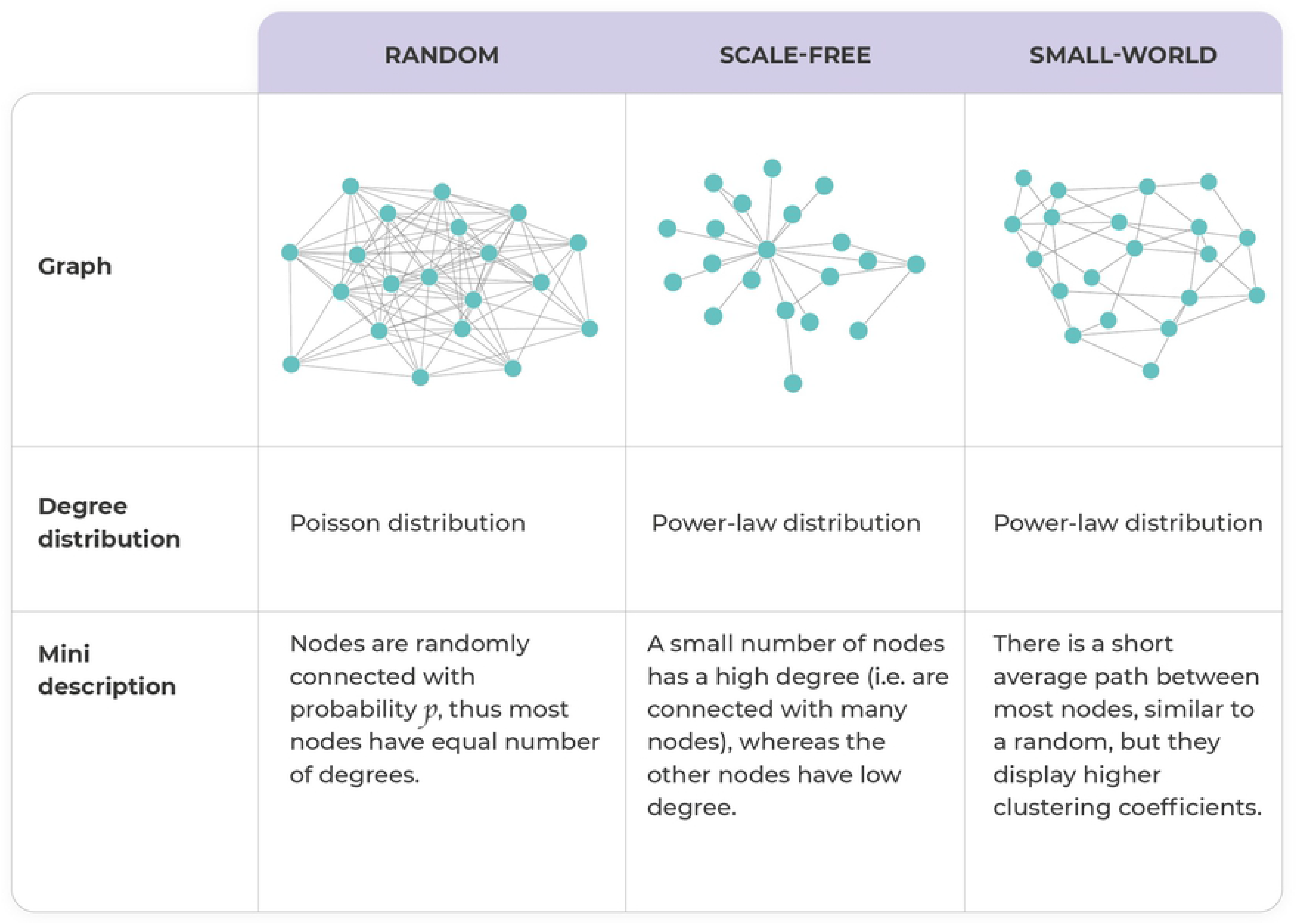
Description of three canonical network topologies. Random, scale-free, and small-world topologies have different network properties. A brief description along with their type of degree distribution in shown.

Degree distributions can also be used to discriminate between network topologies. Thus, we have included in the MicNet toolbox a function that plots the Complementary Cumulative Distribution Function (CCDF) of the degrees of the given network and compares it with the CCDF of a simulated comparable random, scale-free and small-word network on a log-log scale. To calculate the CCDF we first divide the range of the degrees into bins; for each bin we obtain its probability as frequency/total. We used this discrete definition of the Probability Density Function (PDF) to calculate the Cumulative Density function (CDF) as the cumulative sum of the PDF:

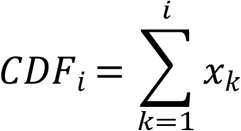

where *x*_*k*_ is the PDF of each bin *k* previously defined, such that the CCDF is calculated as:

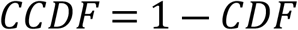

We used CCDF since it has been suggested as an easier way to visualize the difference between degree distributions [84,85].

#### Community Analysis

To analyze subnetworks, we used two ways of dividing the network into subunits: 1) We used Louvain method to detect communities in networks (using python-louvain library [86]), and 2) we used the clusters found with clustering algorithm HDBSCAN. Each subnetwork’s nodes and edges were isolated as a subnetwork using function nx.subgraph, and then for each we obtained the following metrics (also used for network topology analysis): total number of nodes, total number of relationships, density, average degree, and clustering coefficient. Finally, we characterize the diversity of each subnetwork by looking at the total number of different taxa present in each subnetwork at the phylum level.

#### Percolation Analysis

Depending on the network structure, networks can be more or less robust to disruptions. Networks are usually formed by a giant component, which includes between 50% and 90% of the nodes. The formation and dissolution of this giant component is called percolation transition in network theory [87]. The percolation approach consists of removing nodes and their corresponding edges and analyzing how much the network's properties are disrupted [88]. The percolation simulation implemented in MicNet consists of *n* iterations; in each iteration a percentage of the nodes (with default value of 0.1, but this can be specified by the user) is removed along with all of their edges. After removing the nodes and corresponding edges, the following metrics are calculated for the remaining network: density, average degree, number of connected components (this last one calculated using the nx.connected_components function), size of giant component, fraction of nodes belonging to the giant component, the communities found by the python-louvain algorithm and the network modularity. We implemented several percolation approaches: 1) random percolation, in which nodes are removed randomly; 2) centrality percolation, in which nodes are removed by centrality (whether degree, closeness or betweenness centrality), higher values first; and 3) group percolation, where groups of nodes are removed according to a grouping variable provided, such as taxonomic groups or HDBSCAN groups. Consequently, network robustness to different types of disruptions could be assessed by looking at changes in different network metrics.

#### Structural Balance Analysis

Structural balance analysis finds all triangle motifs in the network, that is, nodes that are interacting in triads, and then classifies them as balanced based on the simple analogy that ‘‘my friend’s friend is my friend’’ and ‘‘my friend’s enemy is my enemy’’ [89,90]. This leads to classifying triads of interactions as balanced if they meet this criterion, or as imbalanced otherwise **Fig 3**. A network is considered to be balanced if most triads found in it are balanced. To calculate structural balance, we found all triads in the network using function nx.cycle_basis, and keeping only the cycles of length three. Then, we classified the found triangle motifs into balanced or imbalanced, depending on their mutual correlations. The output of the analysis is a percentage of balanced and imbalanced triangles with respect to all triangles found, and the exact percentage for each of the four types of triangles displayed in **Fig 3**.

**Fig 3.**
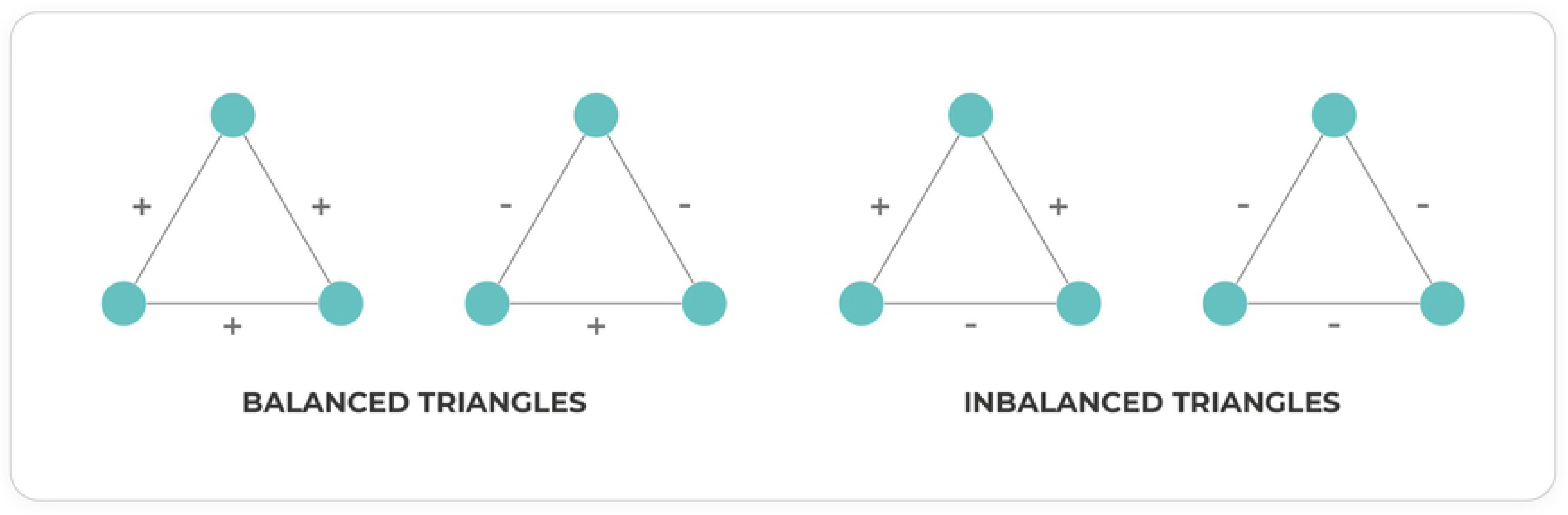
Classification of triad motifs according to structural balance theory. Balanced or imbalanced criterion is assigned according to their mutual correlations.

#### Potential Key Taxa Analysis

Four different centrality measures were implemented to characterize each node (OTU/ASV): degree centrality using function nx.degree_centrality, betweenness centrality using function nx.betweenness_centrality, closeness centrality using function nx.closeness_centrality and PageRank using function nx.pagerank. When running the code, a dataset is returned with each centrality metric for each node.

### Dashboard Interface

In addition to the freely accessible source code at the github repository, we have also developed a web dashboard at http://micnetapplb-1212130533.us-east-1.elb.amazonaws.com that can be used to run most of the analyses presented here. The dashboard consists of three main parts: 1) UMAP and HDBSCAN, 2) SparCC and 3) Network analyses. In the first component, the raw abundance data should be input (see Input Data section), and several parameters, as well as normalizations, for UMAP and HDBSCAN can be modified as desired. This first component returns an interactive visualization of the UMAP plot along with HDBSCAN clusters identified by color. Both the resulting plot and the file detailing cluster belonging can be downloaded from the dashboard.

The second component is for the estimation of the co-occurrence network with SparCC. In this section, abundance data should be input, and parameters can also be adjusted by the user. The resulting correlation matrix can be downloaded from the dashboard. Finally, the third component includes the post-processing analyses of the co-occurrence network. Thus, the input of this section should be the matrix obtained from the previously defined SparCC component or any square correlation matrix, and the UMAP/HDBSCAN output file. When run, this section allows the user to obtain the large-scale metrics of the network, the structural balance percentages, descriptions of the communities found, and two network graphs where the size of the node indicates degree centrality, green edges indicate positive relationships between two nodes, whereas red edges indicate negative ones. Finally, in the graph named HDBSCAN the color of the nodes refers to the HDBSCAN cluster they belong to; whereas in the graph named Community the colors indicate the color of the community they belong to, based on Louvain clustering algorithm. When the network of interest consists of less than 500 nodes an interactive visualization plot is deployed, but for larger networks a static plot is returned, given limited computational resources.

For the other network analyses presented, such as percolation analysis and topology comparison, all the functions necessary to run them are in the github repository and can be used locally by users when downloading the toolbox. It should be noted that, if more computational resources are needed, the dashboard itself can be run after downloading the MicNet toolbox from github, creating the conda environment and deploying it with streamlit as suggested in the readme file.

## Validations

### Modified SparCC and Topology Validations

We performed two validations with simulated communities. We validated the new version of SparCC on the dataset provided by **Friedman & Alm (2012)** [17], which consists of 50 OTUs in 200 samples drawn from a multinomial log-normal distribution. For this, we compared the real correlations to the estimations performed by SparCC, and then we calculated the RMSE (Root Mean Square Error). Secondly, to corroborate topology comparison using large-scale metrics, three networks with known topology interactions were simulated. The networks were constructed with 100 nodes each, and all of them with approximately the same density of ρ = 0.3 and an average degree of 30. Correlation matrices were simulated with three algorithms previously described to obtain: 1) a random network, 2) a scale-free network, and 3) a small-world network. The interaction magnitudes were simulated from a uniform distribution (range from −1 to 1). We calculated the different network metrics on each of these network topologies and used a bootstrap procedure to calculate the distribution of each metric given a certain topology. Finally, we also compare topologies using their degree CCDF, as described in a previous section.

### Biological Validation: Kombucha Consortium

To further authenticate MicNet Toolbox’s approach to analyze microbial co-occurrence networks, we needed to see if biological interactions previously described by experimental work could be replicated in the network. For this, we make use of the kombucha dataset described in **Arıkan et al. (2020)** [91]. Test data was downloaded from the European Nucleotide Archive (ENA) at EMBL-EBI under the accession numbers ERP104502 (https://www.ebi.ac.uk/ena/browser/view/ERP104502) and ERP024546 (https://www.ebi.ac.uk/ena/browser/view/ERP024546). The raw 16S amplicon reads were filtered, processed and annotated with QIIME 2 [92] and DADA2 [93]. Abundance and taxonomy for each ASV cluster was acquired. The obtained abundance table with all samples was filtered, as singleton and unique counts were removed from the data, as suggested by **Berry & Widder (2014)** [25]. Filtering unique and singletons resulted in 48 ASVs. For the visualization module, UMAP parameters were set as follows: number of neighbors of 5, minimum distance of 0.10, number of components of 2 and an Eucledian metric. In the case of HDBSCAN the parameters were: minimum cluster size of 5, minimum sample size of 3 and Bray-Curtis metric. Network construction and network analyses were performed as described in previous sections. The raw data from the kombucha database is in the github repository so that the main results can be replicated and the user could interact with them in the dashboard.

### Case Study: *Archean Domes*

A 16S amplicon dataset was provided from a highly diverse microbial community named *Archean Domes*. This dataset comes from a microbial mat located in the Cuatro Ciénegas Basin (CCB), Coahuila, Mexico (coordinates 26°49’41.7’’ N, 102°01’28.7’’ W). The sampling used in this case study, which consists of ten samples along a 1.5 m transect, represents a natural community with more than 6,000 ASVs [80]. Compositional data was acquired as raw reads form 16S amplicon sequencing. Reads were filtered and processed for clustering and taxonomic annotation in QIIME 2 platform, as shown by the authors [80]. Singletons and unique counts were subsequently filtered as suggested, and consequently, 2,600 ASVs remained [25]. The ASV abundance matrices along with a taxonomic annotation for these sequences were used as input for the MicNet toolbox. For the visualization module, UMAP and HDBSCAN parameters were set as follows: number of neighbors of 15, minimum cluster size of 15, and minimum sample size of 5. Network construction and analyses were implemented as described in previous sections.

## Results

### Validations

#### The Enhanced SparCC

To validate that the modifications performed to SparCC did not affect its performance, we ran our version of SparCC on the dataset provided by **Friedman & Alm (2012)** [17]. We compared our estimated correlation with their true basis correlation (**Fig 4A-B**). We found an overall RMSE of 0.08, and a consistent value of RMSE when estimating small and large correlation values from the simulated samples (**Fig 4C**). Thus, although the original pipeline of SparCC was not modified, by implementing several techniques that parallelized different parts of the code, our implementation of SparCC can now be used for large databases in a reasonable amount of time with relatively small RMSE.

**Fig 4.**
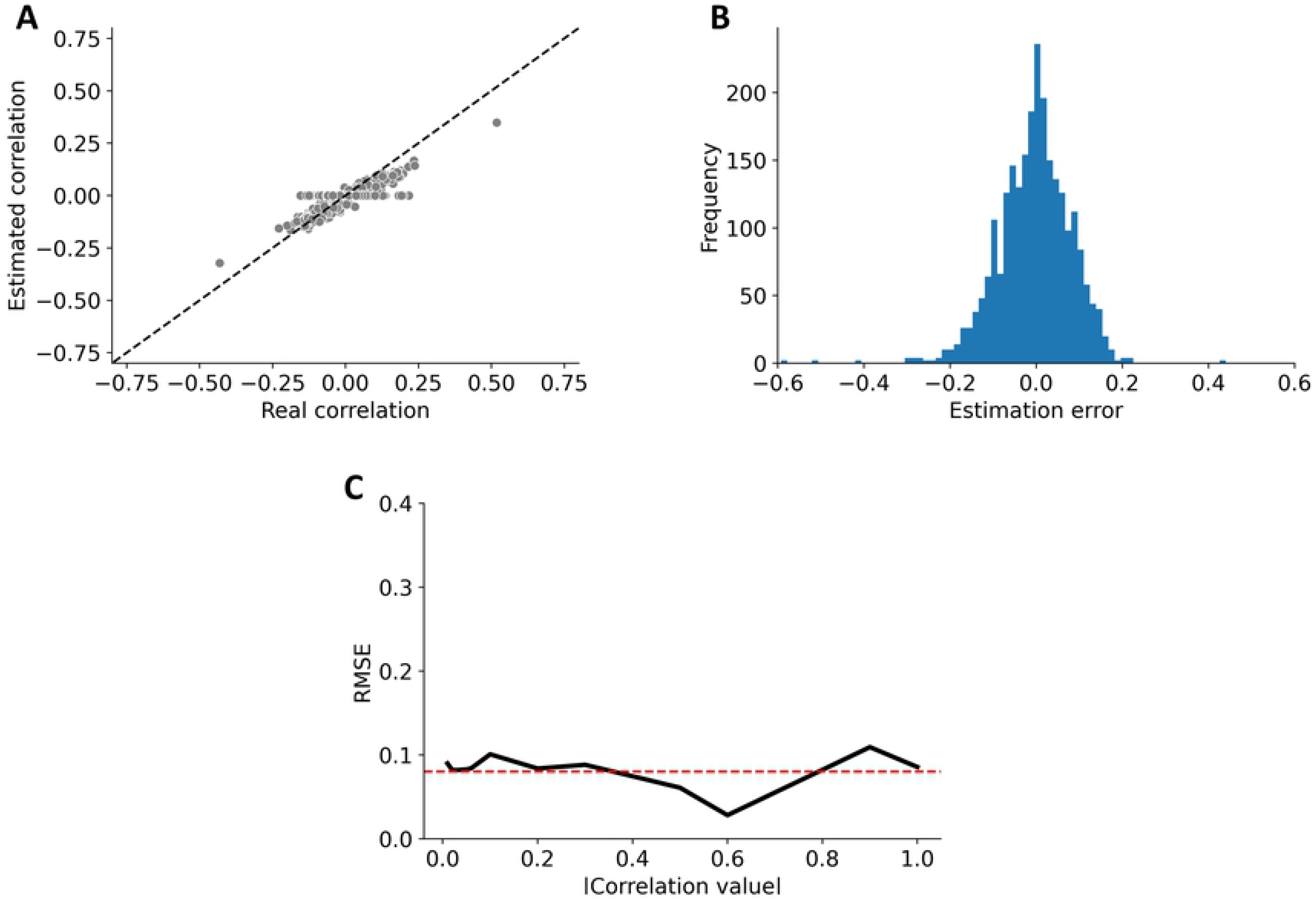
SparCC validation. The modified version of SparCC was validated using the database provided by **Friedman & Alm (2012)** [17]. **A**. Comparison of the estimated correlation with the real correlation. **B**. Histogram of the estimation error produced with the new SparCC version. **C**. RMSE across different absolute values of correlations, the overall RMSE error was 0.08 shown in the dashed red line.

#### Discerning between Network Topologies

One prevailing question when analyzing microbial communities is whether these are randomly assembled or whether they follow certain non-random structures [31,46,94]. In order to validate that large-scale metrics can be used to differentiate between topologies, we began by creating three simulated networks with known random, small world or scale-free topology. All three networks were of 100 nodes each, and all three were made comparable in terms of density and average degree. Then, for each network topology we calculated the following large-scale structure metrics: clustering coefficient, average shortest path length, diameter, modularity, small-world index and degree standard deviation (**Table S1: Supplementary material**). For each topology, we simulated 300 networks and obtained the metrics mentioned above. This allowed us to obtain a distribution of each metric with different underlying topologies.

Results are shown in **Fig 5**, where we can see for each metric three distributions for random, scale-free and small-world networks. It can be appreciated that not all metrics are good to distinguish between topologies. In particular, modularity seems to be good to detect small-world topologies which display a higher modularity than that expected with random or scale-free topologies, but it does not distinguish between random and scale-free networks. Similarly, the network diameter is not a good indicator of topology, in fact, for scale-free and small-world networks, the diameter value in all 300 simulations was always 3, so we were only able to plot a density distribution for random networks. In contrast, clustering coefficient, average path length, degree standard deviation and the small-word index seem to discriminate well between topologies.

**Fig 5.**
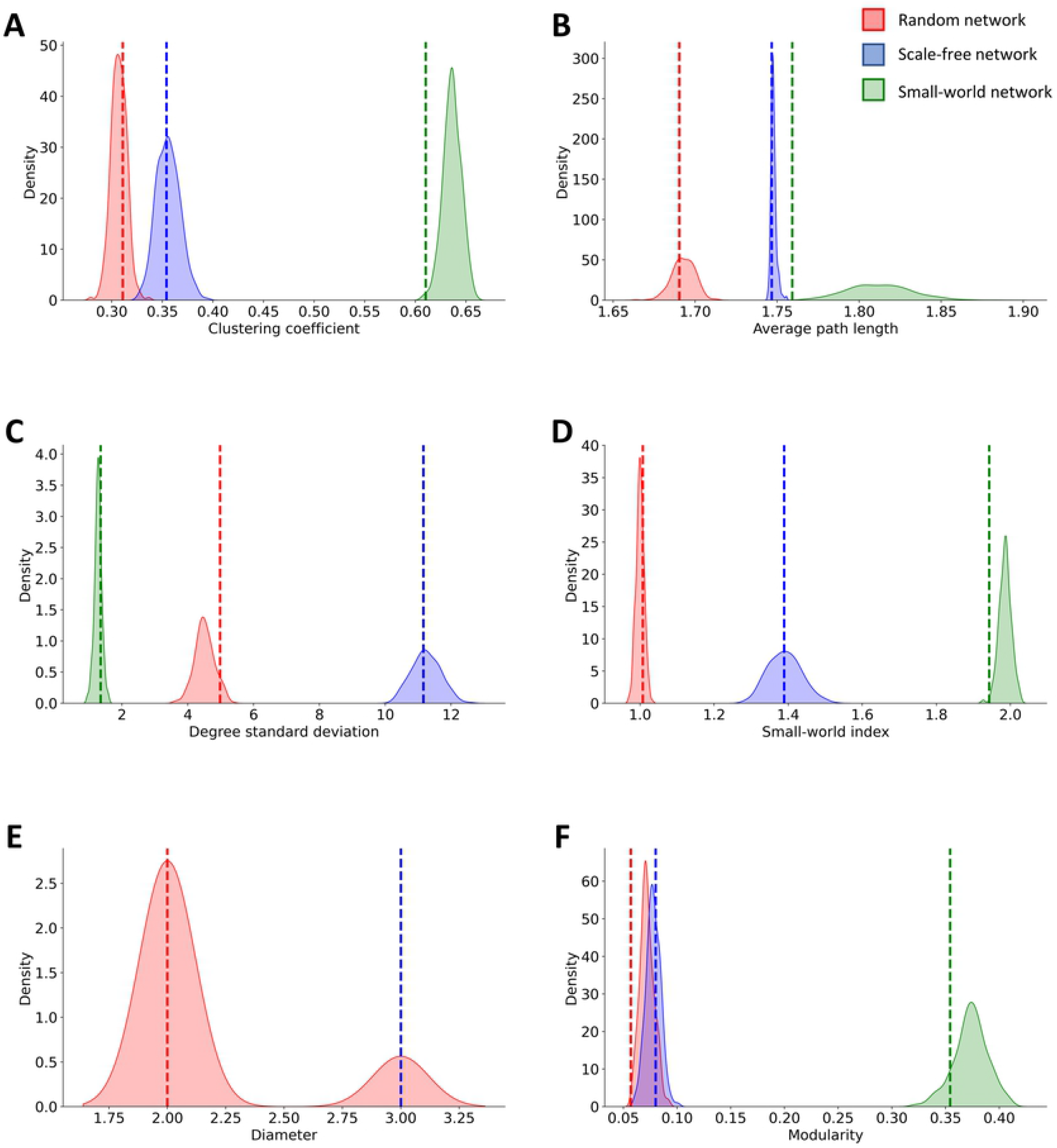
Bootstrap distribution of large-scale metrics according to random, scale-free and small-word topology. Bootstrap distributions for random (red), scale-free (blue) and small-world (green) topologies for the following metrics: **A**. Clustering coefficient, **B**. Average shortest path length, **C**. Degree standard deviation, **D**. Small-world index, **E**. Diameter, and **F**. Modularity. Dotted lines represent the values for each topology shown in **Table S1: Supplementary material**.

Finally, to show how the degree distribution discriminates between topologies, in **Fig 6** shows the histogram of each simulated network topology and the CCDF with a regression line fitted. We can see that the histogram of the scale-free network is clearly different from the other two topologies. Furthermore, the CCDF of the random network had the worst fit, with an explained variance of r2 = 0.66, whereas the small world and scale-free CCDF show an almost perfect linear relationship, with explained variance of 0.89 and 0.88 respectively, as would be expected given their power-law distribution. Thus, degree distributions can also be used to give us hints of the underlying topology.

**Fig 6.**
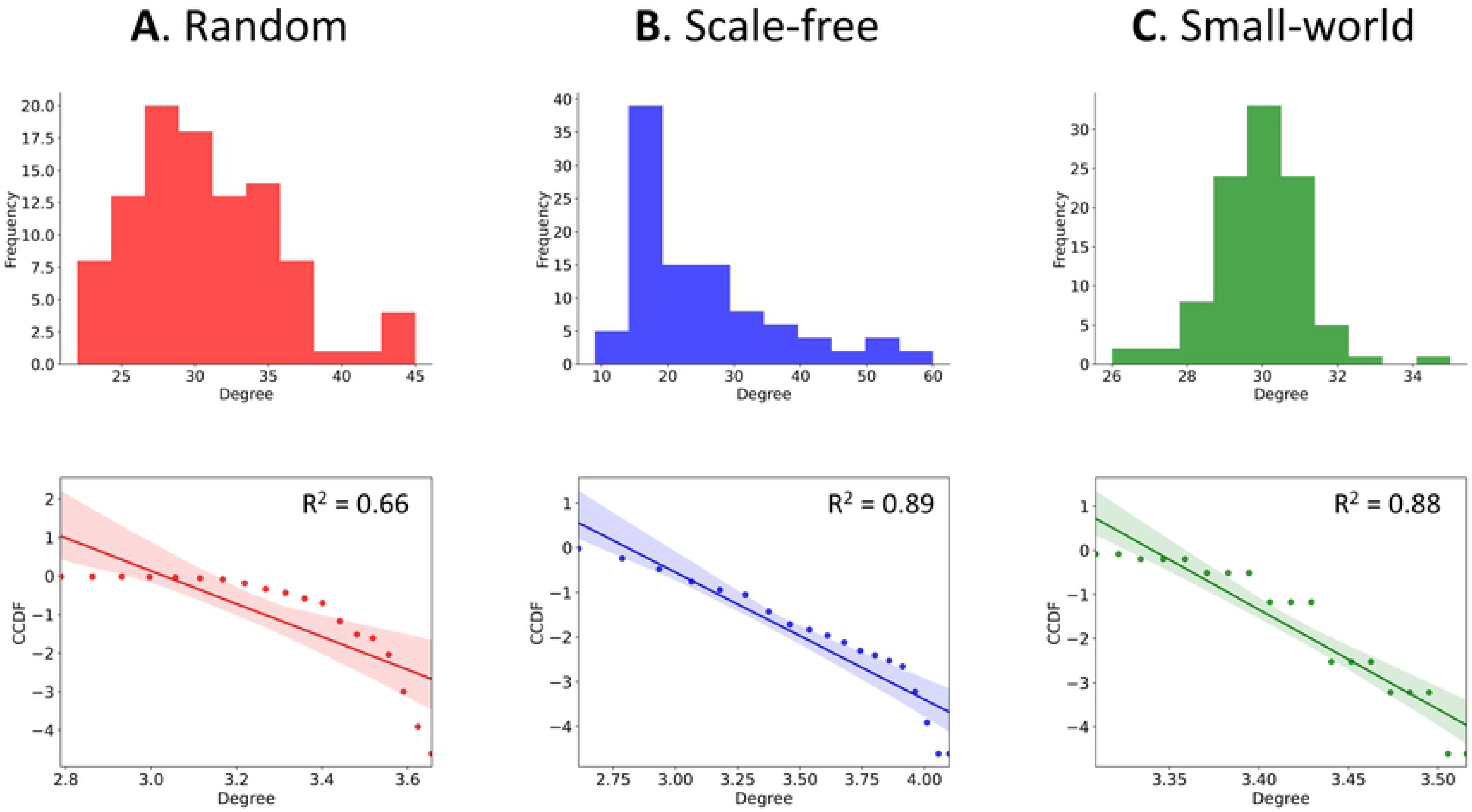
Degree distribution and CCDF for random, scale-free and small-world networks. Another way to differentiate between topologies is with their degree distribution, with CCDF allowing a clearer picture of whether it follows a power-law distribution, characteristic of small-world and scale-free networks. We show the histogram and CCDF with linear regression for an **A**. random network, **B**. scale-free network and **C**. small-world network. CCDF plots are on a log-log scale.

#### Biological Validation: The Kombucha Consortium

To demonstrate how MicNet tools could infer ecological associations, we used a kombucha data set to replicate main global and local behavior. Kombucha is a simple and well-studied microbial consortium of bacteria and yeast, which grow as biofilm due to cellulose production from acetic acid bacteria (AAB) [91,95], but also develops as a liquid consortium. This consortium has been suggested as a convenient tractable system, whose general cooperative and antagonistic multi-species interactions have been previously described [95–97]. After filtering singletons and unique taxa from the raw data, only 48 ASVs remained in the analysis, corresponding to five annotated bacterial taxa and three fungal taxa.

As MicNet pipeline suggests, first, the community was visualized and analyzed with UMAP and HDBSCAN to uncover global patterns and noise taxa. Clusters from HDBSCAN showed one main group containing almost all ASV (34 of 48), a small group with 12 ASV and 2 ASV classified as noise shown in purple (**Fig 7C**). This could refer to a close-interacting community where highly stratified interactions are not common. Since the kombucha community has similar compositions in both homogeneous liquid and biofilms [91], physical closeness between all organisms is expected; this was reflected in the formation of one main group with the HDBSCAN algorithm. We then obtained the co-occurrence matrix of the kombucha samples using our modified version of SparCC. **Table S2: Supplementary material** shows the main metrics of the kombucha network and **Fig 7A,B** show the resulting co-occurrence network.

**Fig 7.**
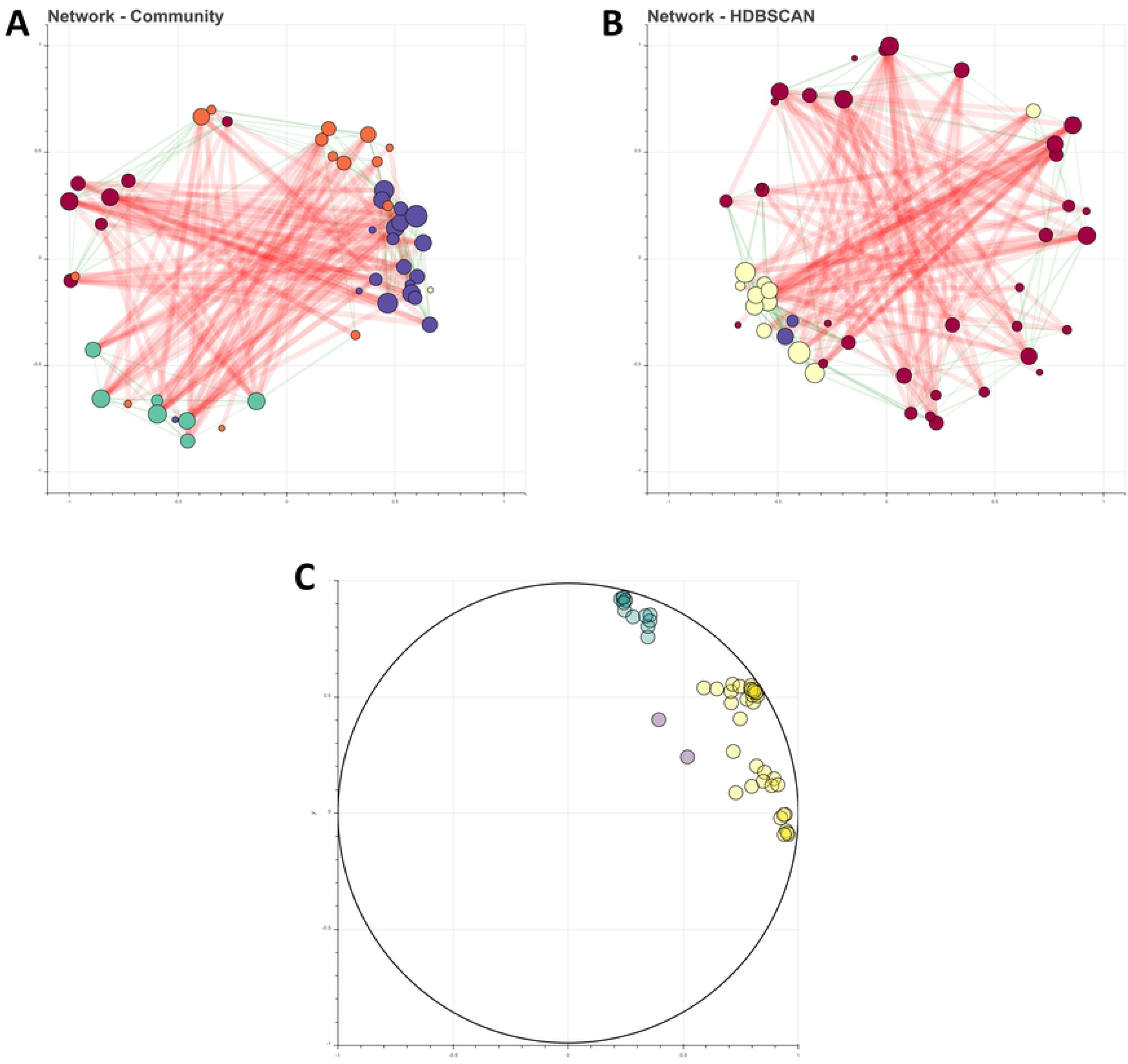
Kombucha microbial network. **A**. SparCC co-occurrence network, where colors indicate Louvain groups. **B**. SparCC co-occurrence network, where color indicates the resulting HDBSCAN clusters. In purple is shown the group depicted as noise. **C**. UMAP and HDBSCAN results show one main group and a smaller one of 12 ASVs.

The resulting co-occurrence network was indeed a relatively connected one, with a connectance of 0.23. This is reflected in the high average degree, which indicates that each ASV is related on average with around 10 ASVs out of the 48 that are present in the community. Highly connected networks could point to a more homogenous environments, including liquid consortiums and slightly stratified biofilms in which kombucha develops, as opposed to more stratified environments, such as soil and microbial mat communities. The kombucha consortium is the result of a metabolic interplay between its microbial consortium, and though cheaters and antagonistic relationships are known [98], more cooperative relationships have been reported [95,96]. In fact, the network did show slightly more positive (55%) than negative relationships (45%), and a major contribution due to mutualist or commensalist interactions is expected.

We perform a topology comparison analysis to explore if the kombucha consortium fits within three canonical networks. Based on metrics bootstrapping from comparable networks, the kombucha network’s degree standard deviation, average path length, SW index suggest some degree of small-worldness [55] (**Fig S2: Supplementary material**). Additionally, comparison of kombucha CCDF with these simulated networks’ CCDF suggest that the degree distribution of kombucha is not following particularly any specific canonical topology, although it appears to fit best with a random network (**Fig S2: Supplementary material**). These results are consistent, as kombucha medium is a non-stratified environment when it develops as liquid consortium, explaining the random properties, but still a stratified microbial consortium when developed as biofilm, explaining the scale-free properties. Thus, kombucha community plausibly shows properties in which microorganisms are adapted to biologically interact with each other in both homogeneous or heterogeneous (to some degree) structures.

Although some biological interactions in the kombucha consortium still need to be confirmed, the global interplay between AAB and yeasts is well-known. In kombucha fermentation, yeast produce invertase which cleave sucrose into glucose and fructose, and further use fructose to produce ethanol. Ethanol is a noxious compound for the consortium. Hence, as a mechanism to regulate ethanol concentrations in the media, AAB transforms glucose and ethanol into gluconic and acetic acid, respectively, exhibiting a straightforward case of syntrophy [91,95]. This biological interplay was depicted in the ASV classified as *Zygosaccharomyces baillii*, the most abundant yeast in the sample, and the ASV with the greatest number of interactions (**Table S3: Supplementary material**). Second to *Z. bailii*, an ASV corresponding to *Komagataeibacter europaeus* (an AAB) appears to be a key central taxa based on each centrality metric. According to the inferred correlations by the enhanced SparCC (correlation matrices at the genus and species level are provided in the Supplementary material as .pkl files), the *Z. bailii* correlated to every other ASVs, and mainly positively co-occurring with *Komagataeibacter* (0.1812), and negatively correlating with *Cortinarius* (−0.2081). Actually, for the species within the *Komagataeibacter* show the highest positively mean correlation with *Z. bailii*, particularly *Komagataeibacter europeus* (0.5397), reflecting the cooperative interplay between yeast and AAB. Additionally, *Komagataeibacter*, an AAB genus, produces acetic acid which inhibits growth of other species, except for *Z. bailii* [91,99]. More specifically, it has been reported that *K. rhaeticus* is one of the main producers of acetic acid compared to other microbial species [100]. As expected, *Komagataeibacter* genus shows negative interactions against some of other species different from their own genus, such as *Variovorax* that is negatively correlated, which might be explained by with its growth-inhibiting capability.

From mean relationships within the taxa present in kombucha, ASVs from the same species tend to have more positive relations between themselves, and this was reflected in the community clustering analysis, where we found that communities were appreciably grouped per species, according to the Louvain method (**Fig S3: Supplementary material**). Clustering resulting from phylogenetic relatedness is common in microbial data, and it may reflect niche overlapping to some degree [4,31]. From the 5 communities predicted with the Louvain method, one of them is considered as noise as it consists of just 1 ASV. HDBSCAN group composition is variable, as most ASV belong to just one group (**Fig S4: Supplementary material**). Nonetheless, the smaller group with 12 ASV’s from HDBSCAN is particularly interesting, as it includes most of *K. europaeus* and the *Z. bailii* ASV, probably depicting the core syntrophic interactions. This potential core group is similarly shown as a Louvain method group. Main metrics for each group of the community analysis (via Louvain or HDBSCAN groups) are reported in **Table S4 and S5: Supplementary material**.

Another aspect of kombucha interactions is that even though yeast could be an important player in the metabolic interplay with AAB, AAB are not fully dependent on them for substrates, characterizing their interaction as some class of non-strict parasitism [97]. In the co-occurrence network, we found evidence that *Z. bailii* was indeed considered a key organism given its high centrality metrics, but in the percolation analysis where nodes were removed by degree centrality (beign Z. bailii), there was not a breakdown of the network (nor in the network density, the average degree, the number of components or the number of communities, as shown in **Table S6: Supplementary material,** along with other percolation analyses performed). In contrast, percolation analysis where the nodes are removed depending on their genus exhibit a network breakup of several components when most of *Komagataeibacter* nodes were removed. Even by removing six *Komagataeibacter* nodes, the network is disrupted into three components, further supporting the relevance of *Komagataeibacter* in the kombucha network. Percolation analysis and centrality metrics are consistent in positioning the *Komagataeibacter* as a crucial genus to the community, and this can be biologically understood due to 1) their independence (to some degree) from yeast to thrive, 2) their cellulose production capability (as a mechanism for protection and resource storage [95]), and 3) as regulators on ethanol concentration.

### Case Study: *Archean Domes* Microbial Mats

To further evaluate the performance of MicNet as a high-throughput toolbox capable of analyzing a highly diverse and complex environment, we tested it on a compositional dataset of ten samples from a microbial mat in Cuatro Ciénegas, Mexico, in the Chihuahuan Desert [80]. This microbial mat, The *Archean Domes*, thrives in a fluctuating hypersaline pond which has been previously described as hyperdiverse [80,101]. Like every microbial mat, it is a stratified community with an intricate metabolic interplay between their organisms [102]. **Espinosa-Asuar et al. (2021)** provided us their microbial data set of 6,063 ASVs, as they have reported. To begin the pipeline, the abundance matrix for all ASV was filtered to exclude low abundance and unique ASVs, remaining 2,600 amplicon sequence variants for the analysis.

First, we search for global patterns and local clustering within the ASV abundance matrix with UMAP and HDBSCAN. The community was grouped into 49 groups, with a mean of 50 nodes in each group (**Fig 8C**). With this approach, the HDBSCAN algorithm allowed us to categorize organisms as noise or outliers, as 553 ASV did not group with any cluster and were categorized as noise, and 260 behaved as outliers.

**Fig 8.**
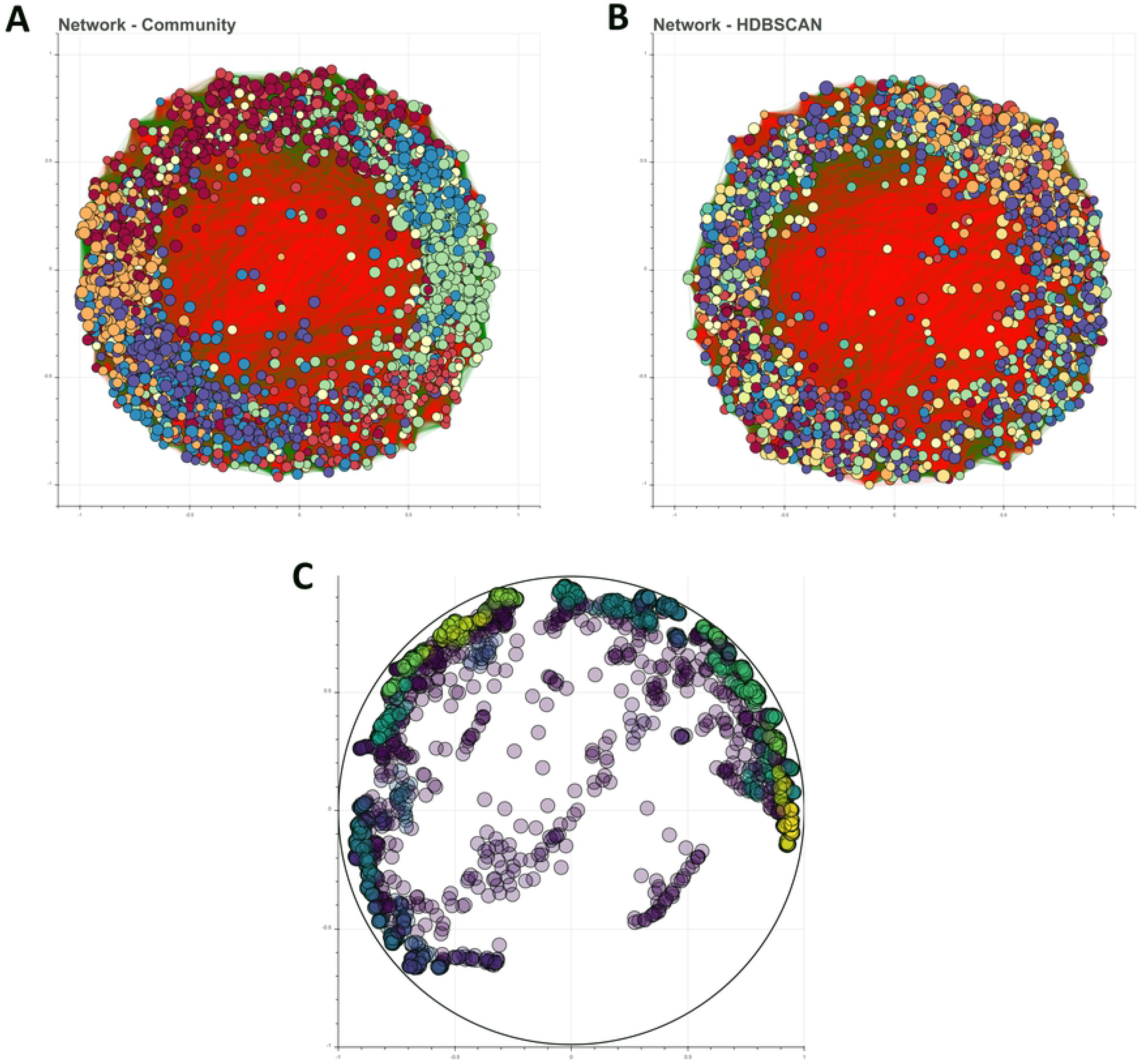
*Archean Domes* biological network. **A**. SparCC co-occurrence network, where different colors indicate Louvain groups. **B**. SparCC co-occurrence network, where color indicates the resulting HDBSCAN clusters. **C**. UMAP and HDBSCAN results show 50 clusters and several ASV classified as noise and outliers

Afterwards, we computed the correlation matrix with the modified version of SparCC. Main large-scale characteristics of the network are shown in **Table S7: Supplementary material**. With the 2,600 ASV we recreated a network with 463,609 interactions with all of them aggregated in a big but sparsely connected network (density = 0.137) (**Fig 8A,B**). This value correlates with the interaction density of an antagonist network between 37 Gammaproteobacteria strains isolated from water samples of *Churince* (reported density = 0.14), a lake situated also in the CCB [103], further suggesting the fact that microbial mats’ networks are often sparse [1,31].

Network topology comparison for the *Archean Domes* network display how high-complexity systems do not fit greatly simplified theoretical models. This was shown in the metrics bootstrapping from simulated comparable networks, where most *Archean Domes* metrics fall in between a scale-free and small-world network distributions (**Fig S5: Supplementary material**). Particularly, degree standard deviation, modularity, SW index, and clustering coefficient suggest that *Archean Domes* possess intermediate properties between scale-free and small-world networks. On the other hand, the average path length of 1.86 is typical of a scale-free network. These results show that this community is not randomly assembled, and that as expected from microbial data, the network is likely an intermediate between a scale-free and a small-world network [31,55].

Potential key taxa analysis based on node centrality was performed for this high-complexity network. An ASV corresponding to a bacterium from the order MSBL9, class Phycisphaerae, phylum Planctomycetes, exhibit the highest centrality values, regardless of the centrality measure employed (**Table S8: Supplementary material**). This bacterial class has been previously described in hypersaline microbial mats as anoxic, fermenting, halotolerant and halophilic microorganisms [104,105]. Looking at other top centrality nodes, most of them were associated to unassigned sequences, except for a node member of the class Parcubacteria and another one from the genus *Imperialibacter* (**Table S8: Supplementary material**). Moreover, unknown taxa as central nodes further suggest the relevance of ‘‘microbial dark matter’’ in ecosystems, and particularly, in hypersaline environments like the one studied here [106].

Local correlations between nodes at different taxonomic levels was inspected (correlation matrices at the phylum and species level are provided in the Supplementary material as .pkl files). From the 463,610 total relationships in the network, we only highlight some of them which might be insightful. At the phylum level, ASVs from Proteobacteria, the most abundant phylum in the sample, have positive mean correlations with most of the phyla, except Nanoarchaeota, Dependentiae and other unassigned prokaryotes. Cyanobacteria, a key phylum in microbial mats, on the other hand face more negative mean correlations with other phyla, including Synergistetes, Acetothermia and Chloroflexi. These negative correlations prospectively originate from overall antagonistic interactions or different niche requirements (such as wavelength, carbon metabolism or temperature adaptation for potential chloroflexi-cyanobacteria associations [107]). Lower taxonomic associations could be inspected. For example, the most central hub taxa (unassigned MSBL9, class Phycisphaerae) positively co-occurs the most (0.5903) with a bacterium from the genus *Dehalobium*, while negatively co-occurs the most with a deltaproteobacteria from the Syntrophobacteraceae (−0.6177).

Although most ecological associations (particularly biological interactions) are analyzed by pairs, species also associate and interact involving more than two organisms, which are vital for ecosystem diversity [108]. Triad motif identification and structural balance theory attempts to address this issue in microbial networks. For the *Archean Domes* network, we identify that most of the triads are balanced, with a highly balanced triad fraction of 0.9995. One property of structural balanced networks is group division, wherein all intergroup ties are negative, and all intragroup ties are positive [109]. Potentially applied to microbial ecology, we suggest that structural balance could reflect niche differentiation, in conjunction with other network metrics and properties that have been previously described as useful [1,4,110].

Furthermore, we carried out a community analysis to inspect subnetwork properties. Our two clustering methods, Louvain and HDBSCAN groups, display contrasting results, which could reflect different ecological structures. Louvain grouping algorithm resulted in 7 network communities with mean total nodes of 371.43 and mean density of 0.4269, while HDBSCAN showed 50 clusters with mean total nodes of 40.94 and mean density of 0.73. Main metrics for each group, for each grouping algorithm, are shown in **Table S9 and S10: Supplementary material**. Phyla composition within most groups (either Louvain or HDBSCAN) is highly diverse, which could mirror spatial structures or compartments at different scales [3,56], which is consistent with the stratified structure of microbial mats [102].

High-diversity communities are commonly associated with an overall stable state. While the high number of ASV in *Archean Domes* probably reflects a highly diverse system, novel methods to assess ecosystem stability have been suggested. In microbiomes, positive relationships alone are prone to destabilize microbial networks, as they can create highly dependent and vulnerable feedback loops [3,111]. *Archean Domes* co-occurrence network show a negative:positive ratio of 0.94, thus, positive and negative relationships are evenly present, suggesting an ecologically resilient, resistant and stablished community. Within this scheme, negative relationships in the community might be originating from antagonistic competitive taxa, lower abundance of facilitative taxa (mutualists), divergent niche requirements, or a combination of all of them [1,3]. Modularity further bolster the stability hypothesis within the microbial mat. Modular groups (or clusters), whether product of biological interactions or habitat preference, plausibly aid in the stability of the system, as fewer links between groups likely ameliorate the spread of local perturbations to other groups. Modularity in *Archean Domes* shows a high value of 0.34, higher than the kombucha network and other published biological networks [3]. Low average path length (1.86 in *Archean Domes* network) likely function as a measure of response capacity to disturbances, hinting about the ecological resilience capabilities of this microbial mat [4,31]. Finally, Percolation analysis delves deeper into community stability. We performed random, centrality and phylum percolation simulations, from which the resulting metrics are shown in **Table S11: Supplementary material**. While none of the node removals induce the breakup of the giant component, the total number of communities and overall modularity do perceive the effects of network perturbations. With this in mind, some degree of ecological robustness could be reflected by the impact of percolation simulations to the co-occurrence network.

## Discussion

MicNet toolbox has shown to be a promising pipeline. Our new implementation of the SparCC algorithm allows larger datasets to be processed without overflowing the RAM in a simple way in our web dashboard. Furthermore, UMAP and HDBSCAN, relatively new dimension reduction and clustering techniques that are promising in microbial ecology studies since, as suggested in this work, they are useful methods to identify metabolic groups, niche overlapping, or subcommunities,. Finally, given the potential of processing co-occurrence networks with a graph theory approach, we have included several network analyses, both new and commonly used, to further describe and understand the resulting networks from SparCC.

One important aspect to have in mind when using the MicNet toolbox is sample size. UMAP and HDBSCAN are known to be quite sensitive to the size of the database used. With a very small dataset (less than 50 OTUs/ASVs) we recommend being cautious at the interpretation level. **Dalmaijer et al. (2020)** [78] have suggested that for optimal use of UMAP and HDBSCAN, it is recommended to have around 20-30 data points per expected cluster or subgroup. Furthermore, the choice of parameters for these two techniques should be done with careful consideration, in particular the number of nearest neighbors and minimum distance [65]. We hope that the interactive dashboard will help in this aspect, since parameters can be modified in a simple way.

In terms of network topology, not all large-scale metrics of a network should be used to discern between topologies. As we show previously, some metrics, such as diameter, have no use to discern topologies, whereas others, like the small-word index provide useful information [55]. Moreover, it is unlikely that any biological system follows exactly a single topology, as shown by our case study: *Archean Domes*, the hyperdiverse microbial mats in CCB. However, we believe that knowing whether a network tends more towards a random, scale-free or small-world could give insightful pointers about its general behavior. For example, a small-world model (created with the Watts-Strogatz algorithm) suggests that the network will have short path lengths, because it is formed by highly connected clusters, which are weakly connected among each other; whereas a scale-free model (created with the Barabasi-Albert algorithm) suggests that networks will have short average path lengths as well, as a consequence that certain nodes that have very high degree and can act as hubs; in both cases the average distance between nodes would be expected to be small but for different reasons, which could give us insight of key biological network properties [85].

Microbial communities in the light of complexity has showed, once again, the potential in drawing biological conclusions from networks. Nonetheless, the biological interpretation of UMAP, HDBSCAN and network analysis should also be taken cautiously. As we mentioned before, the interpretation of the different metrics obtained is debatable, and there is a high diversity of interpretations and terms (see synonyms in Table 1) used for the same ecological concepts. As for today, researchers are encouraged to correlate network theory metrics to biological significance in an attempt to find fundamental metrics that could be useful to describe a particular biological phenomenon in whichever microbial system. With MicNet we suggest an analysis pipeline including visualization, co-occurrence network creation and postprocessing of the resulting network with graph theory analyses that could be used as a standard method for network analysis, offering an overview of a microbial community and enabling the comparison between different microbial systems. Potentially, this approach promises to aid in the search for biologically fundamental metrics.

Biological validation with a kombucha consortium was accomplished, as known local and global behavior, including key taxa and interspecific biological interactions empirically confirmed elsewhere [91,95], were reproduced by the proposed toolbox. Moreover, our high-complexity case study, *Archean Domes*, displays the scope and usefulness of MicNet toolbox by deconstructing microbial co-occurrence networks to manageable biological knowledge. Aware of current caveats of the limitations microbial co-occurrence networks have [1,26], we restate that this approach should be taken as a roadmap for further research on the microbiome system, rather than a conclusive analysis. For example, directed studies to Phycisphaerae bacteria (and other central taxonomic groups) can be performed, and consequently, assess the relevance of this taxon to the whole community structure and functioning. Similarly, ‘‘microbial dark matter’’ characterization and relevance could be further explored with the increasing technologies and databases [112]. Directed co-culture experiments and other novel strategies such as microdroplets are fundamental for biological interactions’ validation [31], which could be applied to inferred correlations between taxa of interest. Moreover, module aimed experiments, including synthetic microbial communities, are tractable strategies that, although reducing complexity of the system, could be informative about mid-scale structures crucial to the system’s stability [113], especially if the experiments include perturbations [114].

Given its potential usefulness, understanding both global and local patterns in microbial communities may be a wise strategy to delve deeper into their currently unknown properties. With the introduction of the MicNet toolbox, we hope that the research community will be able to implement several existing and new analysis techniques in a straightforward manner to further keep unravelling the intricate conundrums that microbiomes hold.

## Availability and Further Directions

MicNet toolbox is available to the scientific community as an open source project at https://github.com/Labevo/MicNetToolbox. In addition, we present an accompanying dashboard which can also be freely visited at http://micnetapplb-1212130533.us-east-1.elb.amazonaws.com, in which the files to build the kombucha network or any other compositional files the user inputs, can be explored and visualized in an interactive way.

## Acknowledgments

We thank Diego Nava for his contributions in the conceptualization and management of the project and Julian Trejo and Diana Fernandez Rosales for their contribution in the elaboration of the artwork. We thank Erika Aguirre for her technical support.

## Supplementary Material

**Fig S1. SparCC parameter selection.** SparCC parameters were set based on our most complex network: *Archean Domes*. We ran SparCC varying **A**. the number of iterations from 10 to 100, **B**. the exclusion number from 10 to 100 and **C**. the exclusion threshold from 0.1 to 0.9. We chose the final values based on the stabilization of the number of edges found, such that the final values used for our databases were: 50 iterations, 10 exclusion number and 0.1 for exclusion threshold.

**Fig S2. Kombucha topology comparison. A**. Distributions obtained from simulated random (red), scale-free (blue) and small-world (green) networks and its comparison to the metrics found in the kombucha network for: degree variance, modularity, average path length, small-world index, and clustering coefficient. **B**. Degree distribution of the kombucha network. We also show the comparison of the kombucha CCDF with a random network CCDF, a small-world network, and a scale-free network.

**Fig S3. Bar plot for community analysis composition (Louvain groups) from the kombucha Network.** Community ID is shown in the x-axis.

**Fig S4. Bar plot for community analysis composition (HDBSCAN groups) from the kombucha Network.** Cluster ID is shown in the x-axis.

**Fig S5. *Archean Domes* topology comparison. A**. Distributions obtained from simulated random (red), scale-free (blue) and small-world (green) networks and its comparison to the metrics found in the *Archean Domes* network for: degree variance, modularity, average path length, small-world index, and clustering coefficient. **B**. Degree distribution of the *Archean Domes* network. We also show the comparison of the kombucha CCDF with a random network CCDF, a small-world network, and a scale-free network.

**Fig S6. Bar plot for community analysis composition (Louvain groups) from the *Archean Domes* Network.** Community ID is shown in the x-axis. Unassigned bacteria and not annotated sequences are grouped in the NA category.

**Fig S7. Bar plot for community analysis composition (HDBSCAN groups) from the *Archean Domes* Network.** Cluster ID is shown in the x-axis. Unassigned bacteria and not annotated sequences are grouped in the NA category.

**Table S1. Network metrics for three canonical topologies**. Large scale metrics of three simulated networks with a random, scale-free and small world topology.

**Table S2. Main metrics and network properties of the kombucha co-occurrence network.**

**Table S3. Centrality measures from potential key players in kombucha network.** Degree, closeness, betweenness, and PageRank centrality was calculated for the top 5 ASV respectively. If a given ASV was not among the top 5 in a centrality measure, the value is reported as NA.

**Table S4. Community analysis (Louvain groups) metrics on kombucha network.** For each community, total nodes, diameter, clustering coefficient, and average shortest path were calculated. Clusters are ordered by increasing density.

**Table S5. Community analysis (HDBSCAN groups) metrics on kombucha network.** For each community, total nodes, diameter, clustering coefficient, and average shortest path were calculated. Clusters are ordered by increasing density.

**Table S6. Network robustness analysis for the kombucha network.** Random, by groups (Genus), and by degree centrality percolation simulations were performed on the Louvain groups. For the percolation by groups, only *Komagataeibacter* and is shown.

**Table S7. Main metrics and network properties of the *Archean Domes* co-occurrence network.**

**Table S8. Centrality measures from potential key players in *Archean Domes* network.** Degree, closeness, betweenness, and PageRank centrality was calculated for the top 10 ASV respectively. If a given ASV was not among the top 10 in a centrality measure, the value is reported as NA.

**Table S9. Community analysis (Louvain groups) metrics on *Archean Domes* network.** For each community, total edges, total nodes, average degree, clustering coefficient, and density were calculated. Clusters are ordered by increasing density.

**Table S10. Community analysis (HDBSCAN groups) metrics on *Archean Domes* network.** For each community, total edges, total nodes, average degree, clustering coefficient, and density were calculated. Clusters are ordered by increasing density.

**Table S11. Network robustness analysis from the *Archean Domes* network.** Random, by groups (Phylum), and by degree centrality percolation simulations were performed on the Louvain groups. For the percolation by groups, only Cyanobacteria percolation is shown.

